# Proteomic Dynamics of Breast Cancers Identifies Potential Therapeutic Protein Targets

**DOI:** 10.1101/2022.06.03.494776

**Authors:** Rui Sun, Yi Zhu, Azin Sayad, Weigang Ge, Augustin Luna, Shuang Liang, Luis Tobalina Segura, Vinodh N. Rajapakse, Chenhuan Yu, Huanhuan Zhang, Jie Fang, Fang Wu, Hui Xie, Julio Saez-Rodriguez, Huazhong Ying, William C. Reinhold, Chris Sander, Yves Pommier, Benjamin G. Neel, Tiannan Guo, Ruedi Aebersold

**Author notes:** Equal contribution (Rui Sun, Yi Zhu, Azin Sayad, Weigang Ge). Co-Correspondence, current address.

## Abstract

Treatment and relevant targets for breast cancer (BC) remain limited, especially for triple-negative BC (TNBC). We quantified the proteomes of 76 human BC cell lines using data independent acquisition (DIA) based proteomics, identifying 6091 proteins. We then established a 24-protein panel distinguishing TNBC from other BC types. Integrating prior multi-omics datasets with the present proteomic results to predict the sensitivity of 90 drugs, we found that proteomics data improved drug sensitivity predictions. The sensitivity of the 90 drugs was mainly associated with cell cytoskeleton, signal transduction and mitochondrial function. We next profiled the proteome changes of nine cell lines (five TNBC cell lines, four non-TNBC cell lines) perturbated by EGFR/AKT/mTOR inhibitors. In the TNBC cell lines, metabolism pathways were dysregulated after EGFR/mTOR inhibitors treatment, while RNA modification and cell cycle pathways were dysregulated after AKT inhibitor treatment. Our study presents a systematic multi-omics and in-depth analysis of the proteome of BC cells. This work aims to aid in prioritization of potential therapeutic targets for TNBC as well as to provide insight into adaptive drug resistance in TNBC.

## Introduction

Breast cancer (BC) is the second most common cancer in the world and the most common in women, with a mortality rate ranking fourth among all malignancies [1]. The major classifications of BCs are luminal A, luminal B, HER2-positive, and triple-negative BC based on membrane receptor expression. Luminal subtypes are characterized by the presence of estrogen receptor (ER) and progesterone receptor (PR), luminal A lacks HER2 while luminal B expresses HER2. HER2-positive BC has an absence of ER and PR receptors, but expresses HER2. Finally, triple-negative BC (TNBC; often referred to as basal BC) lacks ER, PR, as well as HER2 expression. TNBC accounts for 15-20% diagnoses. BCs expressing the aforementioned receptors are usually responsive to drugs disrupting the pathways of the given receptor, whereas the TNBCs currently lack such treatment possibilities. Poly-ADP-ribose polymerase (PARP) inhibitors or immunomodulators, such as antibodies targeting programmed cell death protein 1 (PD-1) and its ligand, are the only therapeutic options for TNBCs. However, the majority of TNBC patients are treated with conventional chemotherapies, and therefore, there is an urgent need for additional treatments and drug targets for TNBC.

Recent genomics-profiling studies have revealed potential novel targets for BC [2, 3]. However, the druggability of these targets remains to be established. As proteins are the predominant catalysts of biochemical reactions, they are also the main therapeutic targets [4, 5]. Therefore, a study of BC proteomics can complement the genomics and transcriptomics studies in drug response prediction and drug target discovery [5]. Cell lines are fundamentally experimental models for drug efficacy tests and simpler model systems than clinical samples or animal models. Several pharmacogenomics cancer cell line databases have been established, such as those from the Cancer Genome Project (CGP) [6], National Cancer Institute (NCI) 60 human tumor cell line anticancer drug screen (NCI-60) [5, 7], and the Cancer cell line encyclopedia (CCLE) [8, 9]. Only few studies have analyzed the proteome of breast cancer cell lines specifically [10–13]. However, they have utilized small numbers of TNBC cell lines. Furthermore, these studies focused unperturbed BC cell lines or tumor specimens collected at a single time point; the response of the BC cell proteome to cancer drug perturbations has not been investigated; finally, the mechanisms of acquired drug resistance were not fully addressed.

Recent advances in data-independent acquisition (DIA)-based high-throughput quantitative proteomic techniques [14] allow high throughput proteome analysis at considerable depth and high degree of reproducibility. We further developed a pressure cycling technology (PCT)-assisted semi-automatic tissue lysis and protein digestion technique to minimize technical variation during sample preparation [15].

In this study, we applied our optimized PCT-DIA method to analyze the proteomes of 76 BC cell lines, including 39 TNBC ones. By integrative analysis of the proteomics results with genomics and transcriptomics data, we generated a model to predict the drug response of these BC cells to 90 drugs and compounds. We then further profiled over 400 BC proteomes from cells perturbed by three drugs prioritized based on our multi-omics modeling of drug response. This unique proteomic data set allowed us to explore the proteins with altered expression patterns in response to the tested drugs and to identify potential drug resistance biomarkers in TNBC cells.

## Results and Discussion

### Proteotyping of 76 Breast Cancer (BC) cell lines

We cultured the 76 BC cell lines characterized in a previous study [2] (**Figure 1A**) Our cell line panel shares 29 lines with the CCLE proteomics study (31 BC cell lines) [8]. Only two lines of the CCLE study are missing in our panel: CAL851 and HCC1500. Furthermore, 45 of 51 cell lines of the Genomics of Drug Sensitivity in Cancer collection [16] were included in our data set. The proteomics data of our 76 BC cell lines were acquired using the optimized PCT-DIA technology [15, 17]. Although other large-scale quantitative proteomics studies have analyzed limited replicates [10–13], the high-throughput of the PCT-DIA technique makes it possible to analyze more replicates and to obtain high-quality data due to its speed. For each cell line, we analyzed four samples, including three biological replicates and a technical replicate of third biological replicate. A total of 302 DIA maps (**Table S1A**) were obtained after quality filtering. Using stringent quantitative criteria as described in the Methods, we consistently quantified 90,762 proteotypic peptides of 6091 SwissProt proteins (**Table S1B**) across all samples with a 34.7% missing rate. For each sample, we identified an average count of around 4000 proteins (**Figure 1B**) and the log10 transformed peptide intensities showed a normal distribution (**Figure 1C**).

**Figure 1.**
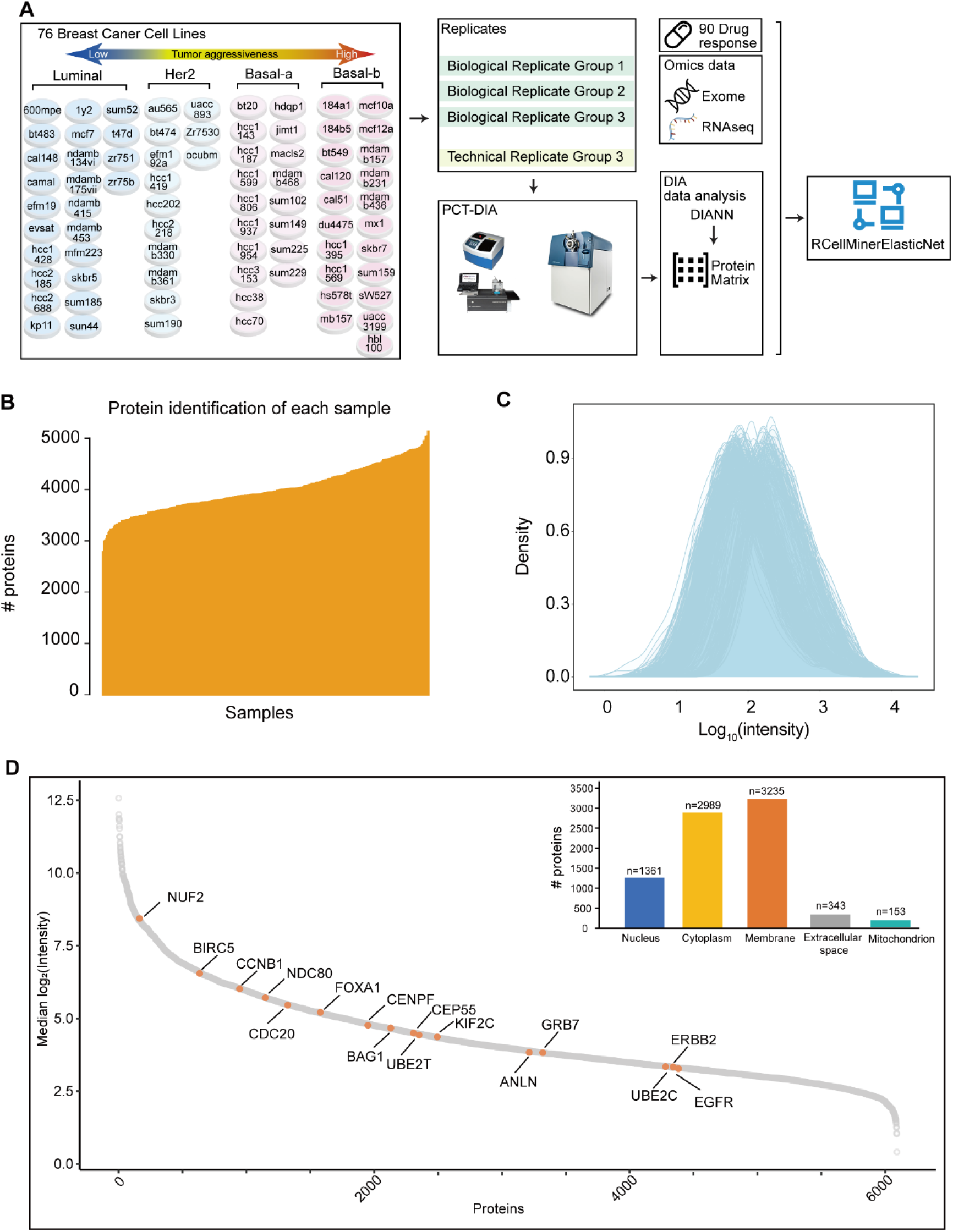
Proteomics analysis of 76 breast cancer cell lines. **(A)** Workflow for generating the proteomics dataset of 76 breast cancer cell lines. The cell lines are classified into four subtypes according to the Neve classification [60]: basal-like A (basal-a), basal-like B (basal-b), luminal, and HER2-enriched (Her2). Four replicates of each cell line were analyzed using PCT-DIA. The proteomics data were integrated with exome and RNAseq data to predict the response of these cell lines to 90 drugs using rcellminerElasticNet. **(B)** Number of identified proteins in each sample. **(C)** Density plot of the quantified protein intensities. **(D)** Identified protein ranking according the median of log2-scaled protein intensity. Orange labeled proteins were overlapped with PAM50.

Replicate proteomic analyses enabled rigorous assessment of quantitative accuracy and batch effects. For each cell line, we analyzed four samples: three biological and one technical replicate (of the third biological sample). Samples from groups 1 to 3 were biological replicates, while samples from the fourth group, which were processed with a different mass spectrometer, were technical replicates of group 3. We observed a higher correlation among technical replicates (median r=0.95) than biological replicates (median r=0.78), which indicates that biological variability was more significant than the technical variability. We next evaluated potential batch effects using principal component analysis (PCA). We observed a mild batch effect of all signal intensities in group 4 only, likely due to it having been analyzed with a different mass spectrometer (**Figure S1A**), as well as using the top 25% highest signal intensities in the four groups to exclude the interference of low abundance proteins (**Figure S1B**). The batch effect was corrected using the limma_R package (**Figure S1C-D**). Within the dataset, we identified 16 proteins overlapping with the PAM50 gene panel, which is a well-known breast cancer classifier (**Figure 1F**). We then computed the average protein intensities of the four groups for subsequent analyses. Comparing the results of RNA-seq (**Table S1D**) [2], reverse phase protein arrays (RPPA) (**Table S1C**) [2], and the present DIA datasets, we found a low correlation between proteomics and transcriptomics data (**Figure 2A**). This was in agreement with previous observations [18]. However, specific genes, such as *ERBB2*, showed a high correlation between their transcription and protein expression levels using both DIA and RPPA detection technologies (**Figure 2B and S2**).

**Figure 2.**
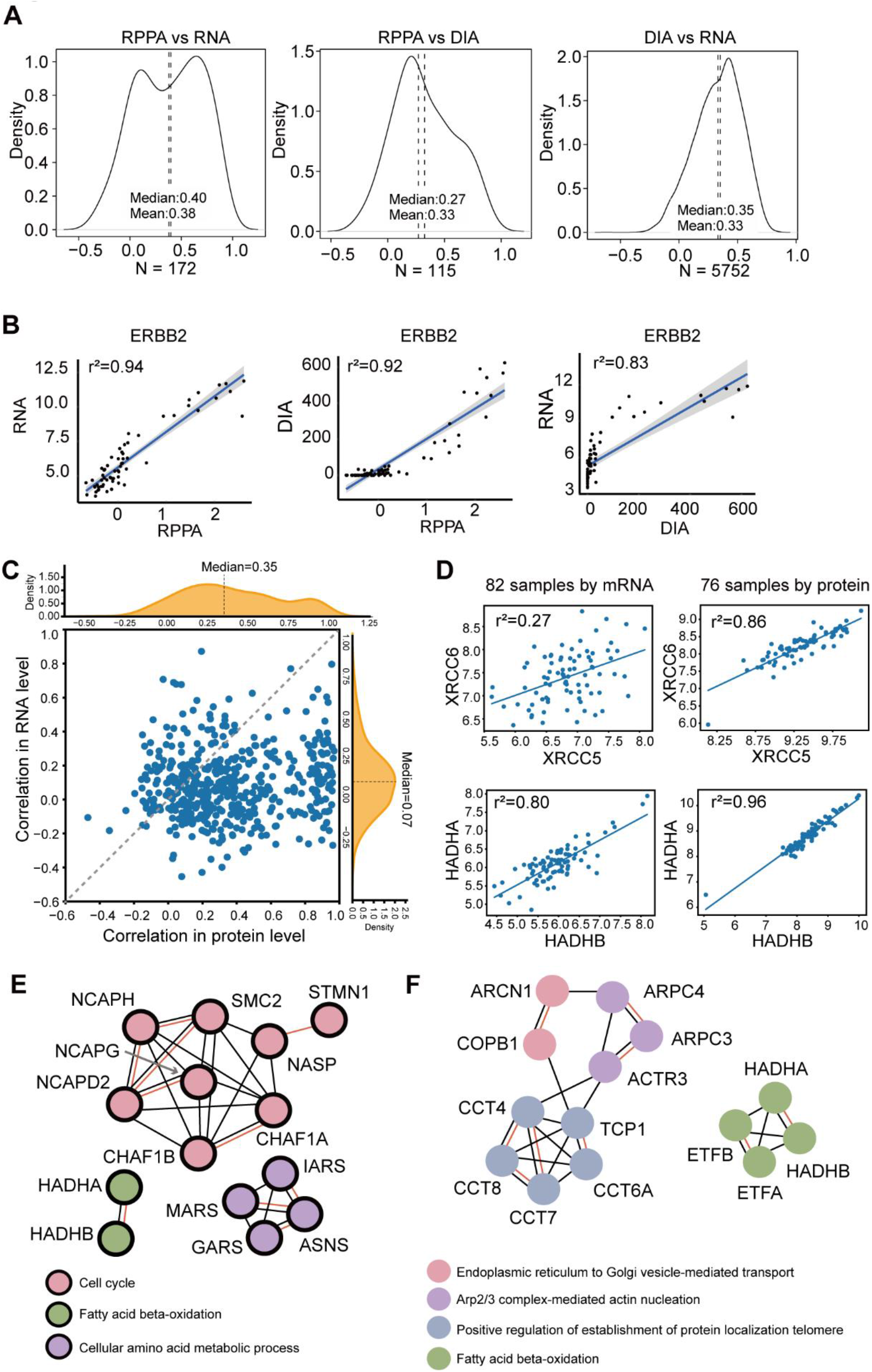
Multi-omics data comparison. **(A)** Density plots of the Pearson correlation coefficients resulting from the pair-wise comparisons between the DIA, the reverse phase protein arrays (RPPA) and the transcriptomics datasets. **(B)** The ERBB2 expression pair-wise comparisons between RNA, RPPA, and DIA, were performed using Pearson correlation. **(C)** Pearson correlation of protein complex between the transcript level and protein level. **(D)** Expression of the protein complexes XRCC6-XRCC5 and HADHA-HADHB at the protein and the transcript levels. **(E)-(F)** Network analyses of the ten protein complexes most highly represented at the transcript **(E)** and the protein **(F)** levels using STRING.

### Detection of 487 protein complexes in BCs

The investigation of protein complexes has been useful for the development of targeted therapies against BC [19]. Therefore, we analyzed the protein complexes of our dataset. Using a reference library containing 622 putative human soluble protein complexes [20], we identified 487 known protein complexes containing at any two subunits in one protein complex. The correlation of the abundance of proteins of complexes (median=0.35) was higher than the correlation of the corresponding transcripts (median=0.07), as indicated by the more remarkable shift to the right shown in **Figure 2C**. This result was consistent with previous reports [21]. We observed that the ten most highly expressed protein complexes showed a higher correlation at the protein level than at the transcription level (**Figure S3**). The cell cycle, fatty acid beta-oxidation, and cellular amino acid metabolic process were the three most significantly enriched pathways among the ten most highly expressed proteins complexes at the transcription level (**Figure 2E**).

While endoplasmic reticulum to Golgi vesicle-mediated transport, actin nucleation, protein localization and fatty acid beta-oxidation were the four most enriched pathways among the ten most highly expressed proteins complexes at the protein level (**Figure 2F**). Notably, fatty acid beta-oxidation was the common pathway detected at both levels of gene expression. The overlapping paired protein complex was HADHA/B (0.96 vs. 0.80 correlation at protein level vs. transcription level) (**Figure 2D**), which is a potential prognostic biomarker for BC [22]. Among the difference in correlation between the protein and transcription level, the proteins from the XRCC6-XRCC5 complex showed a significantly higher correlation at the protein (r=0.86) than at the transcription level (r=0.27) (**Figure 2D**). Their protein complex could bind to DNA and be involved in DNA repair and transcription of selected genes, such as ERBB2 [23]. Similar differences in the correlation of the complex component abundances at protein and transcription level were also apparent in the colon cancer [24]. Therefore, phenotype-related functions may be better reflected at the protein level.

### Identification and validation of 38 proteins differentially expressed in TNBC

To study the distinct molecular patterns in TNBC cells, we found that the DIA data outperformed RNA-seq and RPPA datasets in terms of distinguishing the TNBC samples from 302 samples based on PCA (**Figure 3A**). By comparing the data from TNBC and the non-TNBC cells of the cohort, we identified 771 differentially expressed proteins (DEP)s (**Figure 3B**). Those proteins were enriched in 12 significantly differentially activated pathways using Ingenuity Pathway Analysis (IPA) (**Figure 3C**) and 14 pathways using Representation and quantification Of Module Activities (ROMA) analysis [25] (**Figure S4A**). Both groups of pathways were associated with extracellular matrix, metabolism, and immunity network relationship.

**Figure 3.**
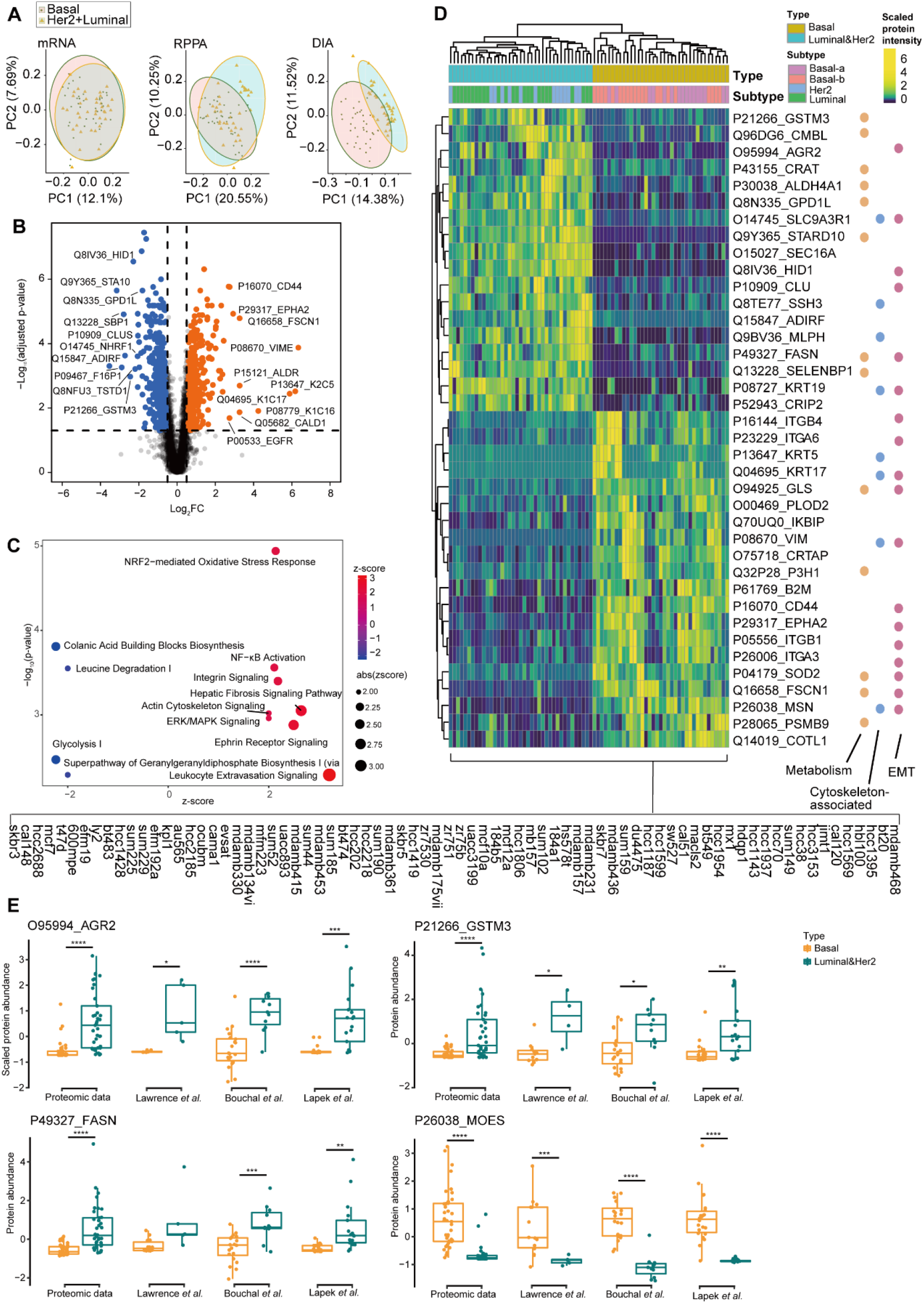
Differentially expressed analysis between TNBC and non-TNBC cell lines. **(A)** PCA plots of the transcriptomics, RPPA, and DIA datasets of TNBC and non-TNBC samples using genes/proteins identified in all three data sets. **(B)** Volcano plot of the differentially expressed proteins between basal and non-basal cell lines (B-H adjusted p-value < 0.05, fold change > 1.5 or < 0.67). **(C)** Bubble plot showing the differentially activated pathways as analyzed by IPA. **(D)** Clustering heatmap of the 76 cell lines showing 38 selected proteins. **(E)** Protein expression of AGR2, GSTM3, FASN, and MOES as measured by our proteomics analysis, as well as by Lawrence *et al*.[35], Bouchal *et al*.[12], and Lapek *et al*.[10].

We next selected the 38 most significantly DEPs to clearly distinguish the TNBC cell lines using unsupervised clustering (**Figure 3D**). These 38 DEPs participated in multiple biological processes (**Figure 3D**) that are known to be involved in BC progression [26, 27]. In particular, 47.4% of the 38 DPEs are the markers of the epithelial-mesenchymal transition [28], indicating that they might contribute to malignant BC. Furthermore, among these proteins, several proteins have been reported to be associated with BC prognoses, such as estrogen-regulated anterior gradient 2 (AGR2) [29], fatty acid synthase (FASN) [30], glutathione S-transferases mu3 (GSTM3) [31] and clusterin (CLU) [32] (**Figure 3E**). Additionally, FASN [30], ITGB4 [33], and ITGA6 [34] have been reported as BC therapeutic targets. We also confirmed the applicability of this panel of 38 proteins for TNBC stratification using unsupervised clustering of three independent breast cancer proteomic data sets [10, 12, 35] (**Figure S4B-D**).

### Proteotyping improves drug response prediction in multi-omics-based modeling

To evaluate whether proteomics improves the multi-omics prediction of a drug response in TNBC and to identify the drug sensitivity biomarkers, we clustered a set of drugs based on the prediction accuracy that emerged from models using various combinations of genomics, transcriptomics, and proteomics data.

We used the rcellminerElasticNet package [5], a wrapper around the glmnet R package and the elastic net algorithm, to explore how integrating various multi-omics datasets affected the response predictions for 90 drugs [2]. To assess the importance of integrating multi-omic datasets, we included in our analysis the following instances: mutations (123 mutations), RNAseq (29,140 mRNAs), proteomics data sets (3672 proteins), and RPPA (218 analytes) (**Figure 4A**). For any drug/data combination, the prediction accuracy was evaluated using the Pearson correlation between the observed and the predicted drug responses (**Table S1C-G**). The predictive power of the DIA proteomic data was comparable to that of the RNA-seq data and superior to that of the gene mutations (**Figure 4B**). The transcriptomic data exhibited a slightly higher correlation than DIA data, probably due to the higher number of transcripts measured compared to the number of proteins, and the fact that we have removed proteins detected only in a subset of the BC cell lines. Similarly, the mutation data had relatively low predictive power that may be attributable to the low numbers of detected mutations, but may also be reflective of the relatively lower levels of mutations found in breast cancer [36].

**Figure 4.**
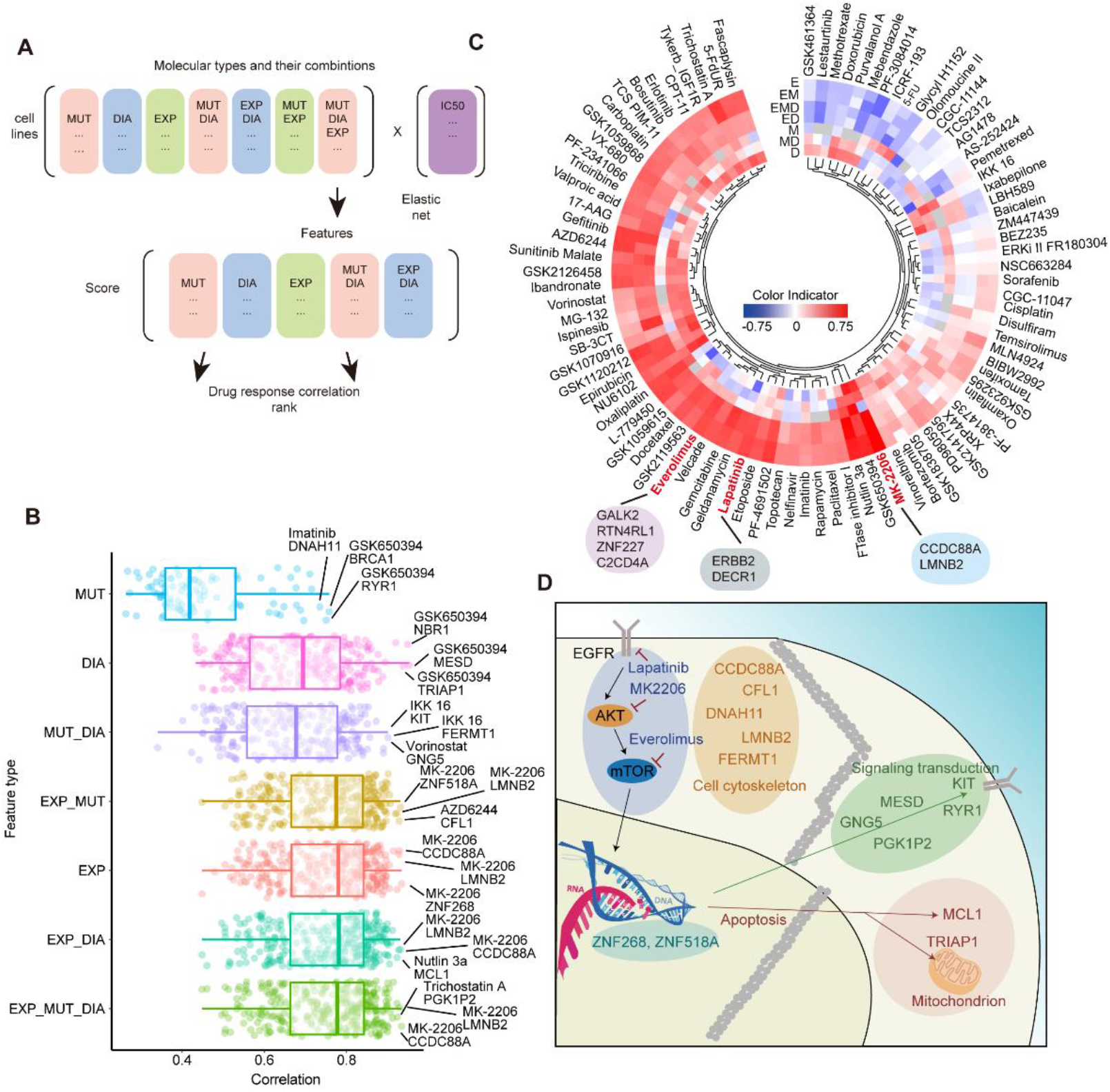
Multi-omics analysis for drug response. **(A)** Workflow of our multi-omics analysis of the drug response by the elastic net algorithm. MUT (M): DNA mutation data (125 mutations); DIA (D) dataset (3672 proteins); EXP (E): transcriptomics dataset (29,140 transcripts). **(B)** Boxplot of drug response predictive power of different dataset combinations (on the y-axis). Each row represents one of the 89 valid elastic net models of the 90 drugs. **(C)** Circos heatmap of the multi-omics results for each drug. Red: drugs for which DIA combined with transcriptomics data showed a better correlation than the transcriptomics data alone would have; the names of three best-predicted drugs are highlighted in red. Blue: transcriptomics data alone provided a better correlation than the DIA and transcriptomics data would have. **(D)** The biological graph showing the protein biomarkers and involving pathways involved in the response to everolimus, lapatinib, and MK-2206 from the elastic net model.

We also displayed the multi-omics drug sensitivity prediction using a circos (**Figure 4C)**; more information is provided in **Table S2**. The various multi-omic data types complement each other in predicting the response to certain drugs. The DIA proteomic data exhibited added value, relative to genomics and transcriptomics data, for multiple drugs, including lapatinib (HER2/EGFR inhibitor), MK-2206 (AKT1/2 inhibitor) and everolimus (mTOR inhibitor). The combination of DIA and transcriptomics data provided better predictions for 21 drugs (**Figure 4C**) than transcriptomics data alone.

We next focused on lapatinib, MK-2206 and everolimus, as the best-predicted targeted therapy drugs. The molecules associated with the response to these three drugs are listed in **Table S2**. To visualize the molecules involved in the response to these drugs, we compiled all the relevant proteins, as annotated by KEGG, into a diagram describing their interactions (**Figure 4D**). These proteins were mainly involved in cell cytoskeleton, signal transduction, and apoptosis, indicating that these biological processes may be associated the drug response to the respective drugs. Particularly, AKT inhibitors (such as GSK650394) can be used to treat BRAC1 deficient BCs [37]. TRIAP1, an anti-apoptosis factor, is a marker of doxorubicin-resistance [38]. As CCDC88A can be phosphorylated by AKT to promote tumor proliferation [39], it might also be influenced by AKT inhibitors, such as MK-2206.

### Protein networks associated with drug response

We next investigated the effect of drugs’ perturbations on the proteome of BC cell lines. In particular, we aimed to identify the protein modules mediating the response to a specific drug treatment and explore the mechanisms for reversing drug resistance. We again focused on the three best predicted targeted therapy drugs, lapatinib, MK-2206, and everolimus. Their effect as explored using nine BC cell lines: SKBR3, T47D, MCF7, Hs578T, ZR75, BT549, MDA-MB-231, MDA-MB-468 and MX-1 (**Figure 5A**). For each drug/cell line combination, we first assessed the half-maximal inhibitory concentration (IC_50_), which measures the amount of a drug required to halve its target biochemical function (**Figure 5B**). Each cell line was then treated with each drug at their specific IC_50_ concentrations, for 4, 12, 24, 48, and 72 hrs; three biological replicates were generated (**Figure 5A**) and lysates of the respective samples were analyzed by DIA MS. A total of 6132 SwissProt proteins were quantified in 432 samples with a 33.6% missing rate (**Figure 5A-C, Table S3A**). The temporal axis of the dataset showed a high degree of reproducibility (**Figure 5D, S5**). In the PCA analysis of each of the three drugs, the luminal cells were well separated from the TNBC cells, indicating that variance of cell types introduce more significant variations than the drug perturbation. (**Figure 5E**).

**Figure 5.**
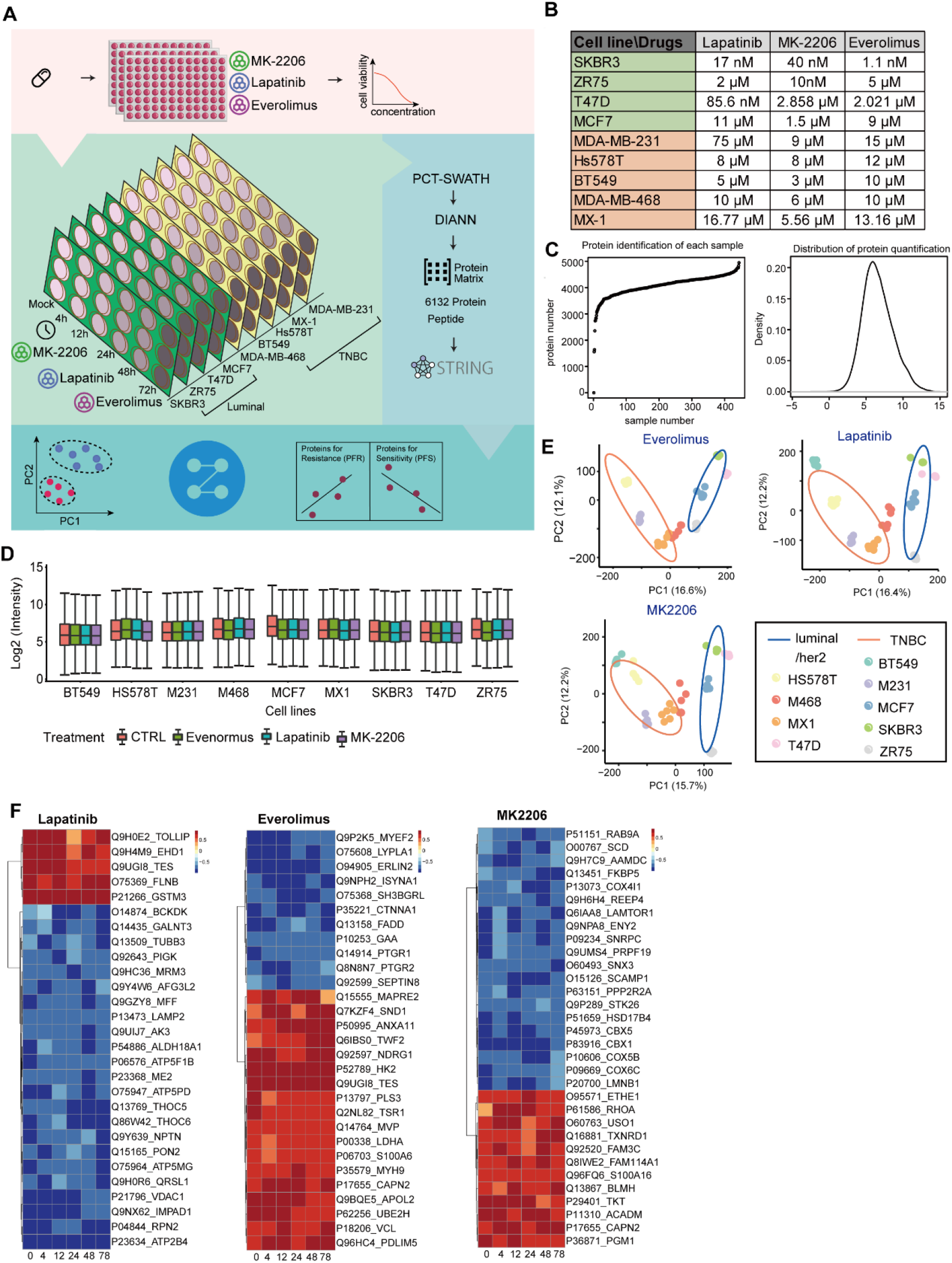
Perturbation experiments on nine BC cell lines using Lapatinib, Everolimus, and MK-2206. **(A)** Workflow of our drug perturbation experiments on a set of BC cell lines. **(B)** The IC_50_ values of each cell line and for each drug. **(C)** The number of identified protein in each sample. **(D)** Density plot of the quantified protein intensities for each cell line. **(E)** PCA plots of the different cell subtypes. **(F)** Heatmaps of correlation values between the protein expression and IC_50_ values for three drugs over time. The x-axis shows the time points (hrs); the y-axis lists the clustered proteins, which included only those with missing values in less than two time points, and mean of correlation coefficient great than 0.7.

We next analyzed the correlation between protein expression and IC_50_ value in each cell line at various time points after drug treatment (**Table S3B-D**). **Figure 5F** shows which proteins positively correlated with resistance (shown in red: 5 proteins for lapatinib, 18 proteins for everolimus and 12 proteins for MK-2206), and which ones negatively correlated with drug sensitivity (shown in blue: 23 proteins for lapatinib, 11 proteins for everolimus and 20 proteins for MK-2206). Among the proteins that positively correlated with drug resistance, several have been reported to be associated with BC therapy. Among them, hexokinase 2 (HK2) can interact with mTOR leading to tamoxifen resistance [40], which is required for tumor initiation and maintenance [41]. CAPN2 which is known to promote castration-resistant prostate cancer invasion via AKT/mTOR [42] has been found upregulated in TNBC patients with respect to HER2 positive BCs [43]. Altogether, these proteins might be associated with the sensitivity of BC after Lapatinib, Everolimus and MK-2206 treatment.

### The dysregulated metabolism of perturbed TNBC cell lines

To compare pattern of proteins that distinguish between the TNBC and non-TNBC cell lines, we used mfuzz and differentially expressed analysis. We identified 252 proteins with the opposite dynamic patterns between the TNBC and non-TNBC cell lines (see Methods, **Figure 6A, Table S3E**). Aligning the protein data with biological pathways the results show that lapatinib and everolimus induced different metabolic responses in the TNBC and the non-TNBC cell lines (**Figure 6A**). This result is in line with previous studies highlighting metabolism as a hallmark of cancer [44]. In particular, our data show that the tricarboxylic acid cycle (TCA), the amino sugar and the nucleotide sugar metabolism, the pentose phosphate pathway, and the glutathione metabolism were upregulated in the TNBC cell lines (**Figure 6A**). However, we did not identify fatty acid oxidation as a predominant difference between the TNBC and non-TNBC cell lines as suggested before [22, 45]. This result suggests that fatty acid oxidation may differ, between the TNBC and non-TNBC cell lines more at baseline, rather than after drug treatment. These metabolic processes we identified occur in mitochondria, which were reported to be dysregulated in the baseline of metastatic TNBC compared with non-metastatic TNBC [46]. Especially, the mTOR signaling pathway may regulate mitochondrial functions [47] Therefore the adaptive changes in metabolism we observed with everolimus may be a consequence of the alteration of mitochondrial function. We observed actin cytoskeleton and mRNA surveillance pathways were altered after MK-2206 treatment (**Figure 6A**). Tumor metastatic require dynamic cytoskeletal activation [48]. The nonsense-mediated mRNA decay, one of mRNA surveillance pathways could induce *BRCA1* mRNA degradation, which is one of the *BRCA1* mutations of breast cancer [49]. These indicated that the cytoskeleton and mRNA surveillance might be important pathways for breast cancer treatment using MK-2206.

**Figure 6.**
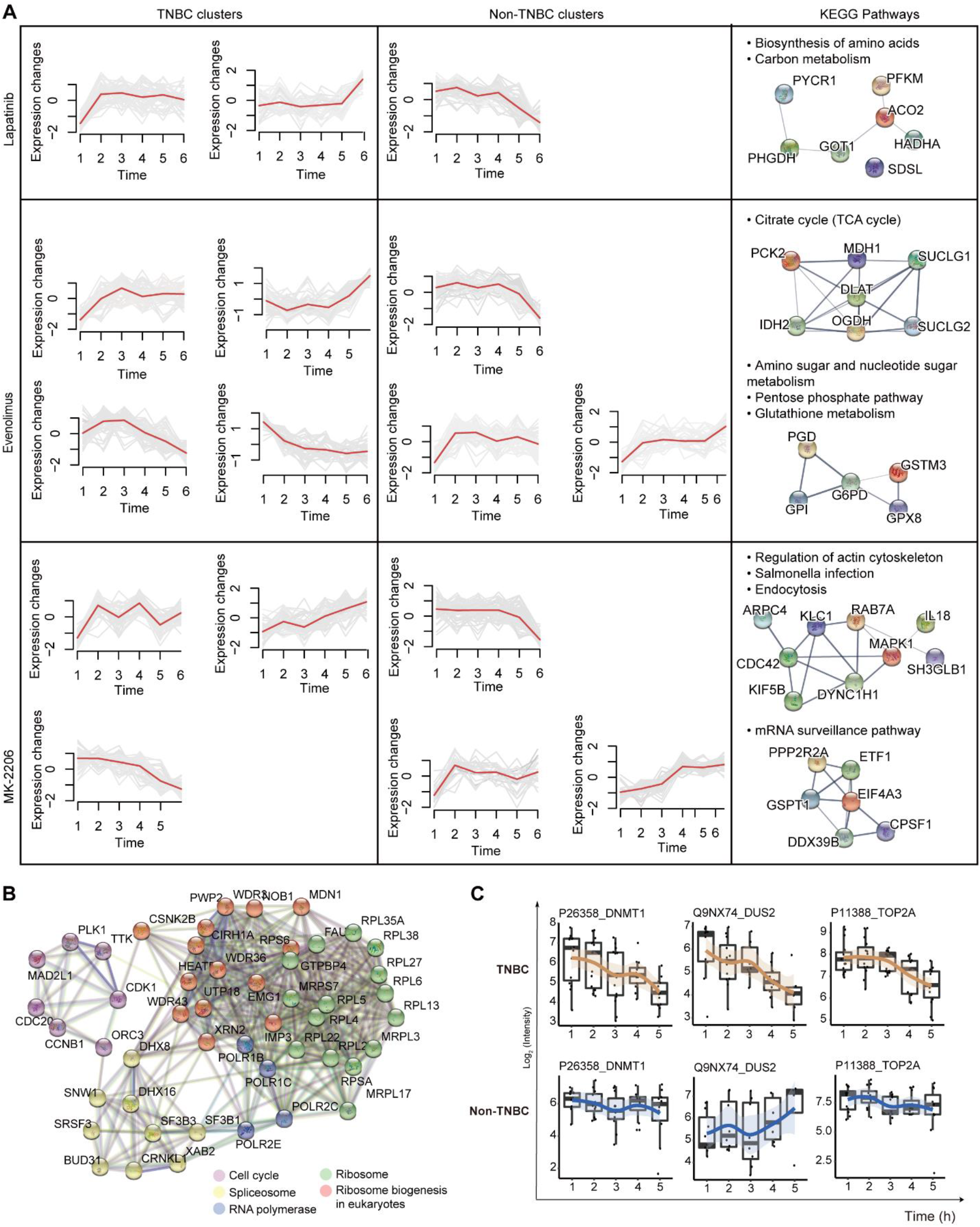
Expression trends comparisons in TNBC and non-TNBC cell lines. **(A)** Time-series of protein expressions showing opposite trends in the NBC and non-TNBC cell lines after treatment with Lapatinib, Everolimus, or MK-2206 using. Left: Mfuzz clustering over six time points (0, 4, 12, 24, 48, and 72 hrs), right: filtered KEGG pathways. **(B)** Network analysis of the DEPs between 0 and 72 hrs in the TNBC cell lines (B-H adjusted p-value < 0.05, fold change > 1.5 or < 0.67). **(C)** Relative abundance of four representative proteins across the TNBC and the non-TNBC cell lines after treated with MK-2206. X-axis: drug treatment time; y-axis: log2-scaled intensity.

To identify the factors that influence the differences in response to lapatinib, MK-2206, everolimus, affecting TNBC and non-TNBC cell lines, we identified DEPs between the pre-treatment and the last post-treatment time point for the three drugs in TNBC and non-TNBC cell lines, respectively. After everolimus treatment, only one protein was downregulated in the non-TNBC cell lines, and none in the TNBC cell lines (**Table S3F**). Lapatinib treatment did not elicit any significant response (no DEP) in the non-TNBC cell lines, whereas seven DEPs were observed in the TNBC cell lines (**Table S3F**). MK-2206 treatment resulted in 182 DEPs in TNBC lines of which only one protein was also identified as DEP in the non-TNBC cell lines (**Table S3F**). The MK-2206 treatment thus induced the most significant difference between the TNBC and non-TNBC cell lines at the proteome level. Specifically, the set of affected proteins mainly included proteins involved in ribosome biogenesis, RNA polymerase, spliceosome, and cell cycle (**Figure S6B**). Among those proteins, we found that DNA (cytosine-5)-methyltransferase-1 (DNMT1) was significantly downregulated only in the TNBC cell lines but no change in the non-TNBC cell lines after MK-2206 treatment (**Figure 6C**). DNMT1 can be activated through AKT phosphorylation to reduce the global levels of DNA methylation [50]. Previous work by others in tamoxifen-resistant breast cancer and colorectal cancers has shown that anti-tumorigenic activities of PI3K/AKT/mTOR inhibitors can be improved by combining DNMT inhibitor [50]. Future study of the current dataset may identify similar drug combination opportunities that exploit adaptive changes in protein level.

Two addition proteins we identified downregulated in the TNBC cell lines but with no change or opposite change in the non-TNBC cell lines after MK-2206 treatment, including tRNA-dihydrouridine (20) synthase [NAD(P)+]-like (DUS2) and DNA topoisomerase 2-alpha (TOP2A) (**Figure 6C**). DUS2 is involved in tRNA modification and affects the stability of tRNA, which could regulate the development of cancer by influencing mitochondrial function [51]. Separately, TOP2A has reported to have key functions in chromosome segregation is a predictor of response for many cancer therapies [5, 52]. We verified that the downregulated expression of TOP2A showed extensive resistance for broad spectrum drugs [53].

Collectively, the proteins and related pathways we identified through analyses of these large dataset lay a foundation for future studies into potential TNBC treatment. Our observations are further reinforced by previous work on the implications of these proteins and pathways in cancer development.

## Conclusion

We collected the high-quality proteomics data of 76 BC cell lines with four replicates using PCT-DIA and integrated these data with genomics and transcriptomics for drug sensitivity prediction. We observed that proteomics data significantly improves the prediction of drug responses in BC. Our analysis identified 38 proteins specifically expressed in TNBC cell lines, which were validated using independent data. Our DIA data set contributed to the effective prediction of drug response in multi-omics modeling experiments. Additionally, we also performed time-course analyses investigating protein dynamics to find which proteins and pathways are specifically dysregulated in the TNBC cell lines and could provide insight to adaptive drug resistance in TNBC. While lapatinib and everolimus elicited few changes in the tested TNBC cells, MK-2206 resulted in significant changes including ones involving RNA modification and cell cycle pathways. Our data showed significant response with DUS2, DNMT1, and TOP2A to MK-2206 in the TNBC cells. These observations warrant future investigations to explore their suitability as drug targets or biomarkers in TNBC.

## Limitations

Although this multi-omics study included a relatively large number of samples, as well as biological and technical replicates, it did not include phosphorylation proteomics data. However, this limitation dose not compromise the relevance and accuracy of our main findings. The primary focus of this publication is to report on this rich proteomic dataset and conduct initial, exploratory bioinformatic analysis of the dataset. It is outside the scope of the intended work, and therefore a limitation, to conduct in-depth experimental validation of reported biomarkers; this work is left for future study.

## Materials and Methods

### PCT-DIA analysis of the cell lines

PCT-DIA analysis was performed as previously described [54]. The fresh cell lysates were processed according to a previously published protocol [15]. All samples were spiked with iRT peptides (Biognosys, Schlieren, CH) [55]. A total of 1.5 μg of cleaned peptides was analyzed by LC-MS/MS as previously described [56]. The ion accumulation time for the MS1 and MS2 acquisition was set to 150 ms and 30 ms for, respectively. The DIA window schemes were optimized to 66 variable windows. The instrument was operated in a high sensitivity mode.

### Elastic net analysis

We used a flexible network algorithm to create a multivariate linear model to predict the effect of each compound in 76 BC cell lines based on genomics, transcriptomics, and proteomics datasets, as previously described [5].

### DIA data analysis

The DIA data were processed using the DIA-NN (1.7.15) and a cell line library [15]. Briefly, DIA raw data files were converted in profile mode to mzML using msconvert. The parameters were left to their default values and the files were analyzed using DIA-NN, as previously described [57].

### Cell culture

BT549, Hs578T, ZR75-1, and MDA-MB-231 cell lines were cultured in DMEM/F-12 medium (Biological Industries, Cromwell, CT, USA); MDA-MB-468 was maintained in L15 medium (Biological Industries); T47D was adapted in DMEM medium (Biological Industries); MCF7, MX-1, and SK-BR-3 were kept in 1640 medium (Biological Industries) supplemented with 10% fetal bovine serum (Biological Industries) and penicillin-streptomycin (HyClone) at 37°C with 5% CO_2_.

### Cell proliferation assay

IC_50_ was determined using the MTT cell proliferation assay. The assay was performed using the manufacturer’s recommendations [58]. Briefly, 5,000 growth phase cells were plated in each well of a 96-well plate, with 100 μL medium. After 24 hrs, the drug gradient concentration was replaced, including lapatinib (Targetmol, 231277-92-2), AKT1-2 inhibitor MK-2206 (Targetmol, 1032350-13-2), and everolimus (Targetmol, 159351-69-6). The cells were then incubated for 72 hrs. Untreated cells were used as negative controls. While the wells without cells were used as blank controls. Next, 20 μL of the 5 mg/mL MTT reagent (Sigma) was added to each well, and the plates were incubated for 4 hrs at 37°C. After incubation, the remaining MTT solution was removed and 100 μL DMSO (Biofroxx, Germany) were added to each well to dissolve the purple formazan and lyse the cell to release the mitochondrial residues of formazan. Absorbance measurements were performed at 570 nm, and the IC_50_ values (i.e., the concentrations that inhibit cell proliferation by 50%) were obtained using Multiscan Spectrum (BioTek, the USA).

### Drug Treatment

Cells were exposed to different drugs, at their IC_50_ concentrations during 4, 12, 24, 48, and 72 hrs. They were then washed with PBS three times before cell collection using a 2000 rpm centrifuge for 5 min. After discarding the waste, dry pellets were preserved at −80°C.

### Opposite dynamic proteins selection

The proteins were clustered in TNBC and non-TNBC cell lines using Mfuzz [59]. We only considered those proteins that: (i) consistently up- or down-regulated over time (B-H adjusted ANOVA p-value < 0.05) with every drug; (ii) showed a fold-change between the last and the first time points higher than 2. We then searched for pathways including these proteins using KEGG pathways in STRING. Pathways consistently dysregulated in at least three out of five TNBC cell lines, and at least two out of four non-TNBC cell lines were prioritized.

## Data and materials availability

The 76 breast cancer cell DIA data are deposited in Pride (PXD004701). The DIA data of cell lines drug perturbation are deposited in iProX (IPX0001825001). All the data will be publicly released upon publication.

## Author contributions

R.A., T.G., and B.N designed and supervised the project. Y.Z., A.S., R.S., W.G., S.L. conducted proteomic analysis. C.Y., J.F. F.W., H.X., and H.Y. contributed to cell culture and drug treatment. A.L., L.S., V.R., J.R., H.Y., W.R., C.S., Y.P. contributed to data analysis. R.A., T.G., R.S., A.L., and Y.Z. wrote the manuscript with inputs from co-authors.

## Acknowledgments

The work was supported by the SystemsX.ch project PhosphoNetX PPM (to R.A.), the Swiss National Science Foundation (grant no. 3100A0-688 107679 to R.A.), the European Research Council (grants no. ERC-2008-AdG 233226 and ERC-20140AdG 670821 to R.A.), the Westlake Startup Grant (to T.G.), Zhejiang Provincial Natural Science Foundation of China (Grant No. LR19C050001 to T.G.), Hangzhou Agriculture and Society Advancement Program (Grant No. 20190101A04 to T.G.), National Natural Science Foundation of China (General Program) (Grant No. 81972492 to T.G.), National Science Fund for Young Scholars (Grant No. 21904107). Part of this study was supported by the Center for Cancer Research, National Cancer Institute, National Institutes of Health (Z01-BC 006150 to Y.P., A.L. V.N.R and W.C.R). We thank Dr Luxi Zhang for assistance for the sample preparation and Mr Guan Ruan, Zhicheng Wu, and Qiushi Zhang and Luis Tobalina Segura for assistance for the data analysis of this manuscript.

## Declaration of interests

T.G. and Y.Z. are shareholders of Westlake Omics Inc. W.G. are employees of Westlake Omics Inc. The remaining authors declare no competing interests.

## Supplementary figure legends

**Figure S1.**
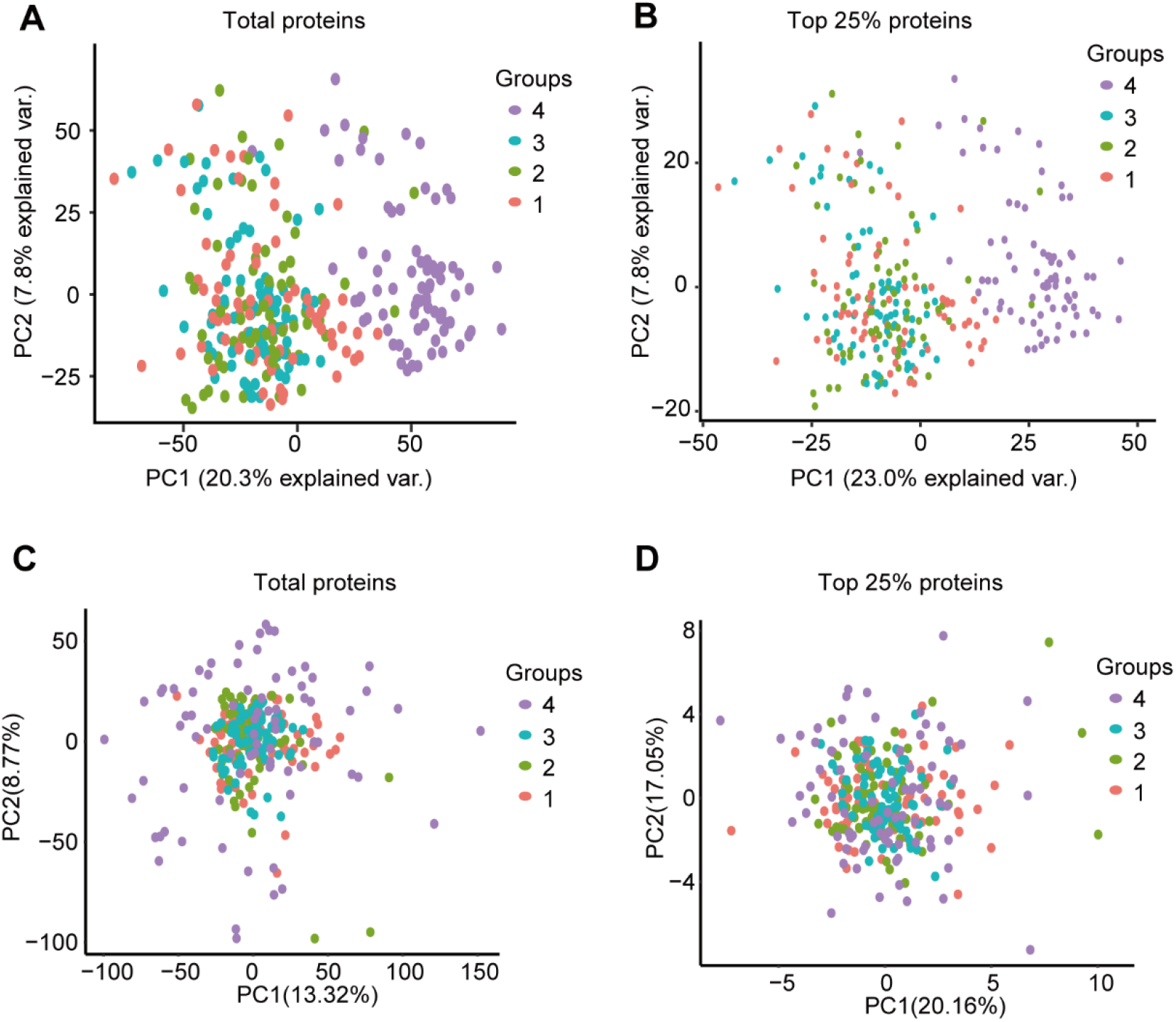
Quality control analysis of the proteome data of 76 breast cancer cell lines. **(A)-(B)** PCA plots of all samples among four groups using total 6091 proteins **(A)** or the 25% most abundant ones **(B). (C)-(D)** PCA plots of all samples among four groups using total all 6091 proteins **(C)** or the 25% most abundant ones **(D)** after batch effect removal.

**Figure S2.**
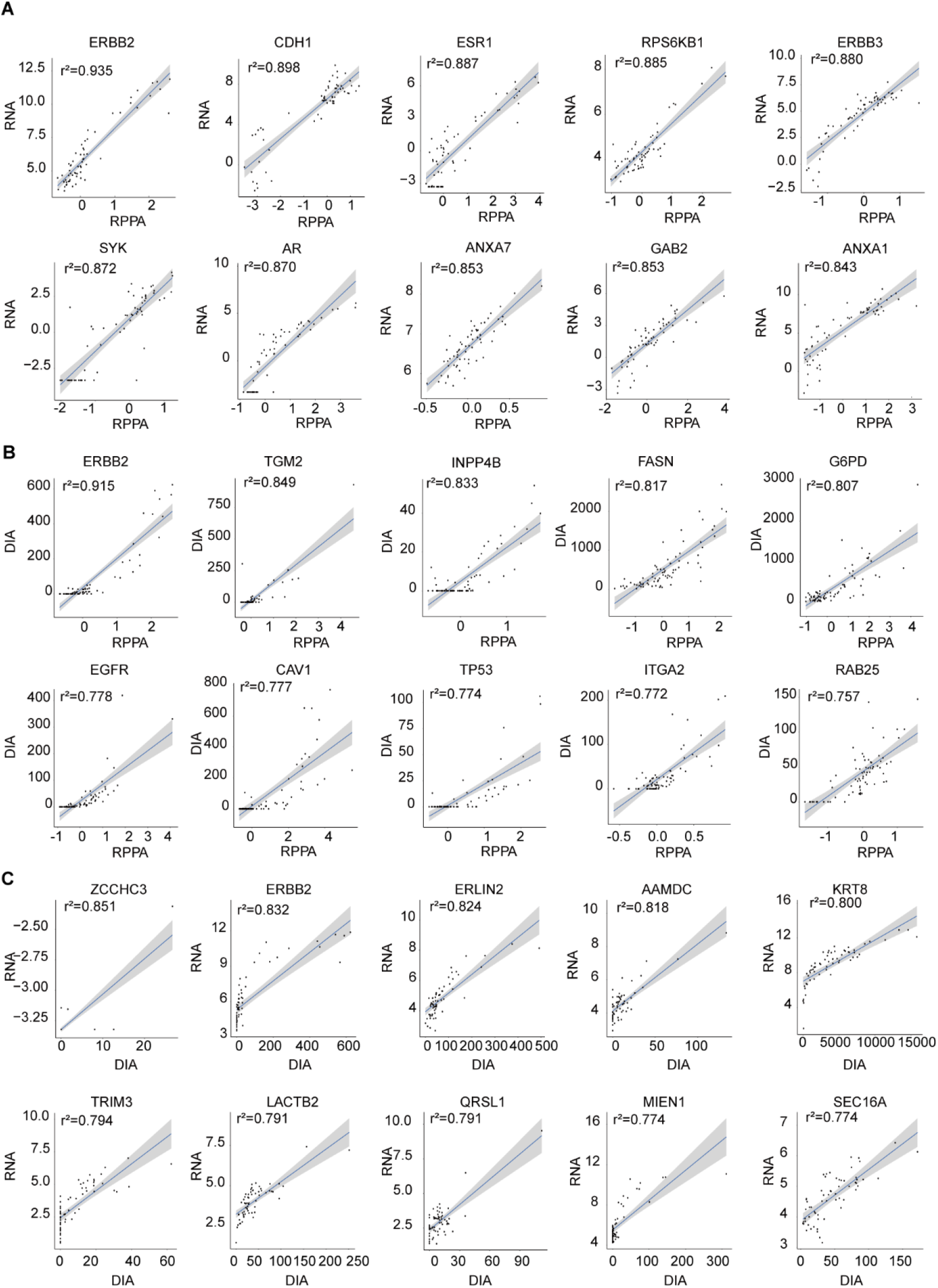
Correlation between mRNA and protein data for selected individual proteins. **(A)-(C)** Expression correlations of ten proteins between mRNA and RPPA **(A)**, RPPA and DIA **(B),** and mRNA and DIA **(C).**

**Figure S3.**
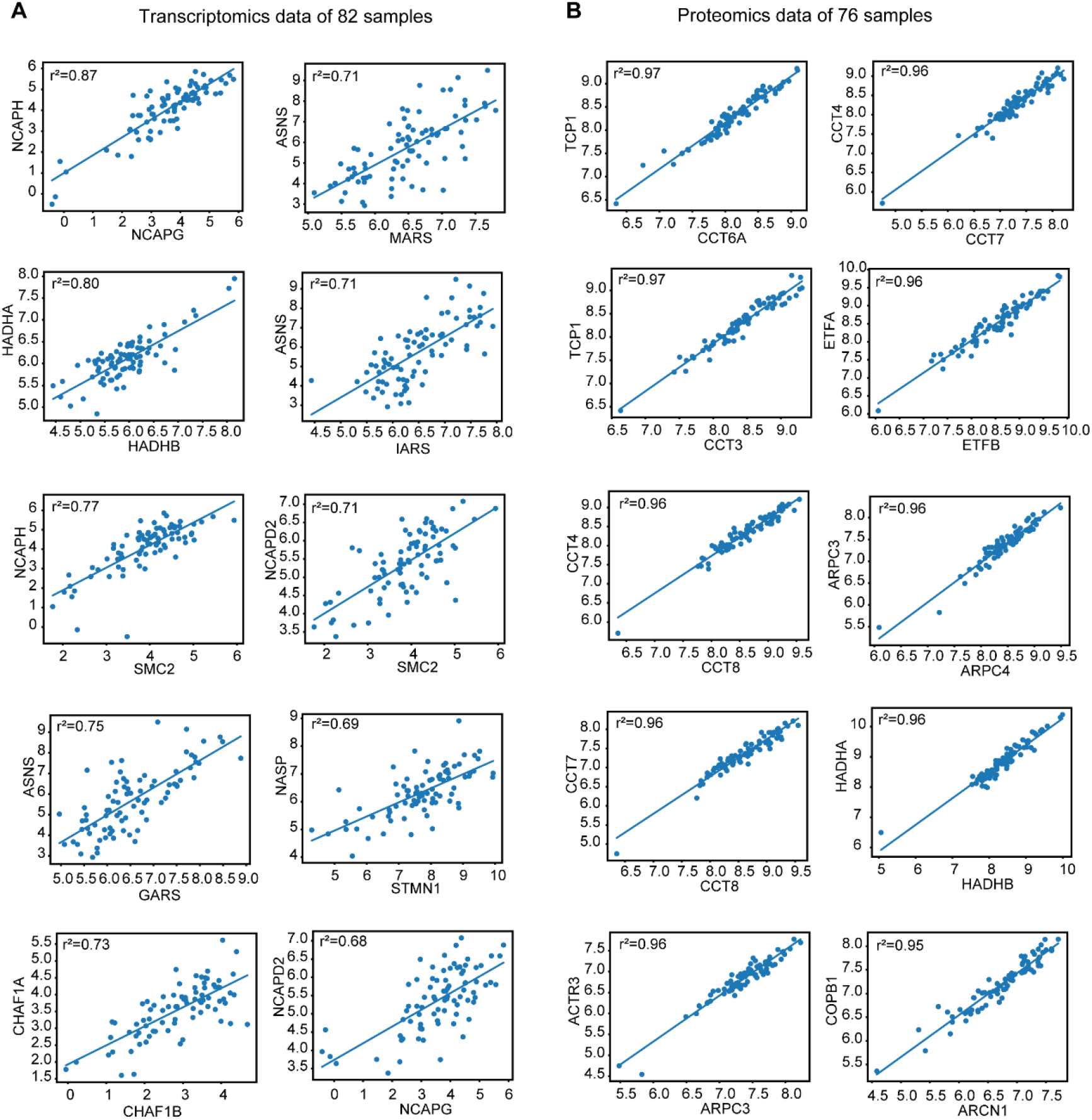
Correlation between mRNA and protein data for the components of selected protein complexes. **A)-(B)** The 10 most correlated protein complexes at transcript **(E)** and the protein **(F)** level using STRING.

**Figure S4.**
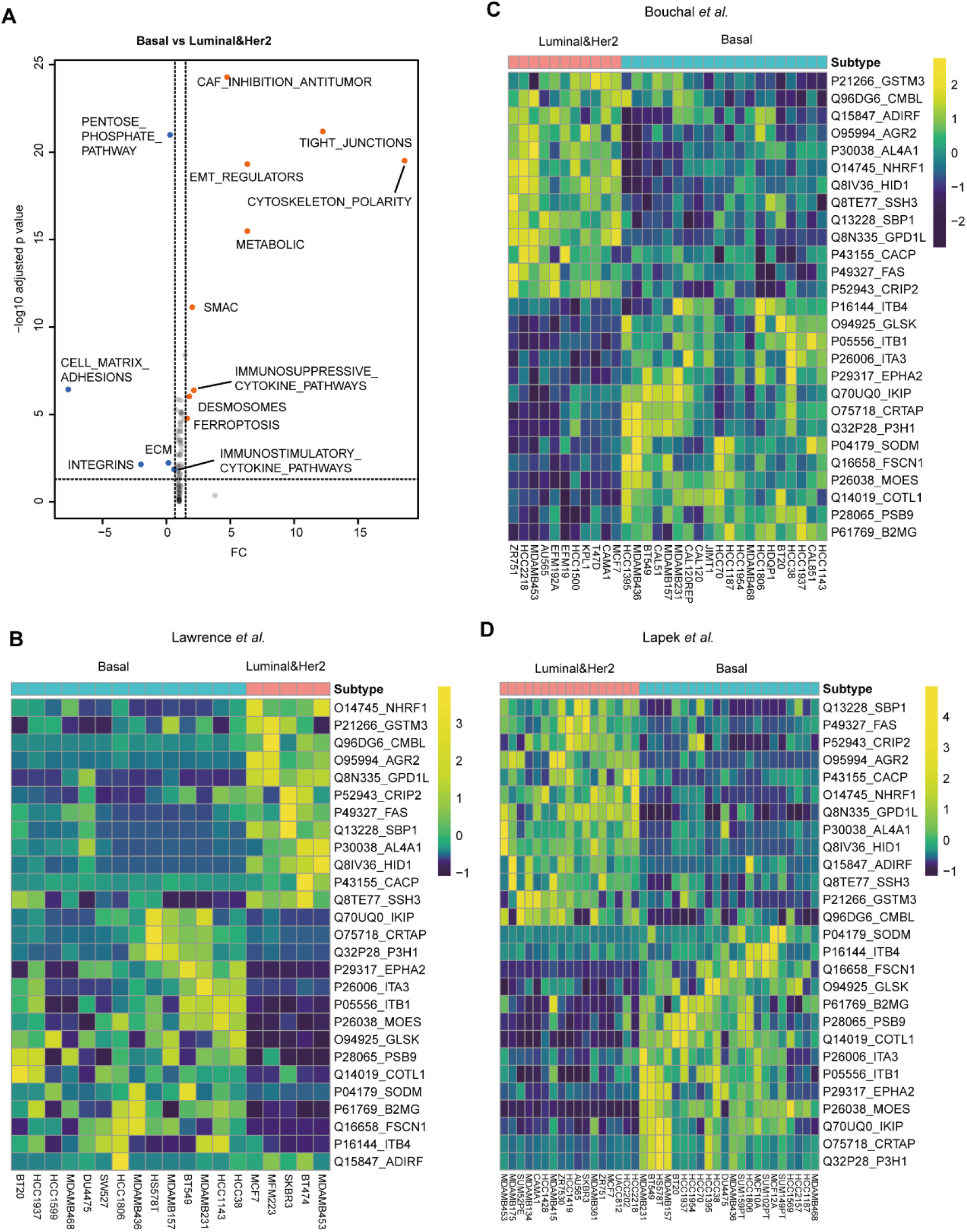
Validation of the 38-protein signatures in four independent datasets capable of distinguishing the TNBC and the non-TNBC cell lines. **(A)** ROMA analysis highlighting the fourteen pathways that best stratify the breast cancer cell lines. **(B)** Validation of the 38-protein signatures using the Lawrence *et al*. dataset [35]. **(C)** Validation of the 38-protein signatures using the Bouchal *et al*. dataset [12]. **(D)** Validation of the 38-protein signatures using the Lapek *et al*. dataset [10].

**Figure S5.**
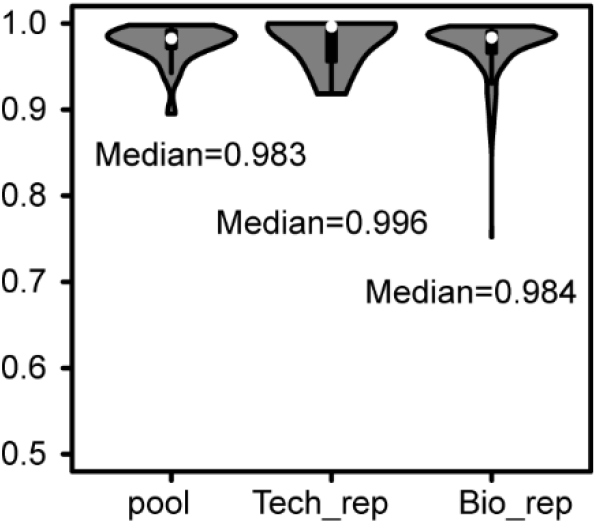
Quality control analysis of the drug perturbation experiment of over nine BC cell lines. The number of identified protein.

